# Deep immune profiling of the maternal-fetal interface with mild SARS-CoV-2 infection

**DOI:** 10.1101/2021.08.23.457408

**Authors:** Suhas Sureshchandra, Michael Z Zulu, Brianna Doratt, Allen Jankeel, Delia Tifrea, Robert Edwards, Monica Rincon, Nicole E. Marshall, Ilhem Messaoudi

**Author notes:** Corresponding Author: Ilhem Messaoudi, Molecular Biology and Biochemistry, University of California Irvine, 2400 Biological Sciences III, Irvine, CA 92697.

## Abstract

Pregnant women are an at-risk group for severe COVID-19, though the majority experience mild/asymptomatic disease. Although severe COVID-19 has been shown to be associated with immune activation at the maternal-fetal interface even in the absence of active viral replication, the immune response to asymptomatic/mild COVID-19 remains unknown. Here, we assessed immunological adaptations in both blood and term decidua from 9 SARS-exposed pregnant women with asymptomatic/mild disease and 15 pregnant SARS-naive women. In addition to selective loss of tissue-resident decidual macrophages, we report attenuation of antigen presentation and type I IFN signaling but upregulation of inflammatory cytokines and chemokines in blood monocyte derived decidual macrophages. On the other hand, infection was associated with remodeling of the T cell compartment with increased frequencies of activated CD69+ tissue-resident T cells and decreased abundance of Tregs. Interestingly, frequencies of cytotoxic CD4 and CD8 T cells increased only in the blood, while CD8 effector memory T cells were expanded in the decidua. In contrast to decidual macrophages, signatures of type I IFN signaling were increased in decidual T cells. Finally, T cell receptor diversity was significantly reduced with infection in both compartments, albeit to a much greater extent in the blood. The resulting aberrant immune activation in the placenta, even with asymptomatic disease may alter the exquisitely sensitive developing fetal immune system, leading to long-term adverse outcomes for offspring.

## INTRODUCTION

The current pandemic of coronavirus disease 2019 (COVID-19) caused by severe acute respiratory syndrome coronavirus 2 (SARS-CoV-2) continues to be a significant threat to human health globally (1). Overwhelming evidence suggests that pregnant women are a high-risk group for COVID-19 (2). Indeed, a study on severe COVID-19 infections in pregnant women from 18 countries have reported higher rates of adverse outcomes such as mortality, preeclampsia, and preterm birth (19). While pregnant women with a severe COVID-19 diagnosis are at 62% higher odds of getting admitted to the ICU compared to non-pregnant women of reproductive age (2, 3), a majority of those who get exposed to the virus experience asymptomatic or mild version of the disease (4-7). Both vulnerability and immune responses to viral infections during pregnancy can be distinct compared to non-gravid individuals, as observed in influenza, hepatitis E, varicella, and measles (8-10). These differences are primarily driven by peripheral immune adaptations during pregnancy (11, 12) that balance fetal tolerance and growth with host defense. Recent analysis of the peripheral immune system of mothers with asymptomatic disease revealed an increase in low-density neutrophils (LDN) but no gross changes in leukocyte frequencies, activation, and function (13). Furthermore, the cytokine storm that is characteristic of severe COVID-19 in the general population is hardly reported in cases of severe COVID-19 amongst pregnant patients (14, 15). These findings support the hypothesis that pregnancy limits the induction of exuberant peripheral inflammatory responses to SARS-CoV-2 infection that are widely reported in non-pregnant individuals.

In addition to changes in blood, the immunological landscape of the maternal-fetal interface (placenta) undergoes significant changes during pregnancy. The decidual compartments of placenta harbor maternal immune cells (macrophages, NK cells, and T cells (16)), which exhibit mixed signatures of activation and regulatory phenotype that correlate with gestation (18) and can respond to foreign particles at the maternal fetal interface (17). However, details of decidual adaptations to a respiratory infection such as COVID-19 are slowly beginning to emerge. Although available data strongly suggest a lack of vertical transmission (5, 21-23) with rare detection of viral RNA in the placenta (24), severe COVID-19 has been shown to trigger maternal inflammatory responses at the maternal-fetal interface (20, 21). Specifically, an increase in markers associated with preeclampsia and activation of placental NK, T cells and increased expression of interferon-related genes and stress associated heat shock proteins have been reported (20). However, placental immune rewiring with mild/asymptomatic infections and how that relates to peripheral immune adaptations remains poorly understood.

Therefore, in this study, we examined the effect of maternal, asymptomatic SARS-CoV-2 infection occurring during late pregnancy on peripheral and decidual immune cells of the maternal-fetal interface at term. We used single cell RNA-sequencing and multicolor flow cytometry to profile changes in immune landscape of maternal decidua. Our analysis revealed an expansion of CD69+ T cells and selective loss of HLA-DR^high^ regulatory macrophages and regulatory T cells (Tregs) with asymptomatic/mild SARS-CoV-2 infection in the decidua. Single cell analysis revealed broad activation of myeloid cells in the decidua, with enrichment of cytokine secreting subsets, attenuation of interferon signaling and a concomitant drop in expression of MHC class II molecules in blood monocyte derived macrophage subsets. Activated T cells and terminally differentiated CD8 T cells (TEMRA) were enriched with infection, but expansion of cytotoxic CD4 and antiviral CD8 T cells was only observed in blood. While cytokine signaling modules were attenuated in the blood, elevated interferon signaling signatures persisted in the decidua. These findings highlight that even mild/asymptomatic infection can trigger placental immune activation with potential long-term consequences for the developing fetus.

## MATERIALS AND METHODS

### Study Participants and Experimental Design

This study was approved by University of California Irvine Institutional Review Boards (Protocol ID HS# 2012-8716) and Oregon and Health Sciences University (eIRB# 22713 and 16328). Informed consent was obtained from all enrolled subjects. All participants in this study were healthy and did not report any significant comorbidities or complications with pregnancy. Detailed characteristics of participants and experimental breakdown by sample are provided in Supp. Table 1.

### Blood processing

Whole blood samples were collected in EDTA vacutainer tubes. PBMC and plasma samples were isolated after whole blood centrifugation 1200 g for 10 minutes at room temperature in SepMate tubes (STEMCELL Technologies). Plasma was stored at -80^0^C until analysis. PBMC were cryo-preserved using 10% DMSO/FBS and Mr. Frosty Freezing containers (Thermo Fisher Scientific) at -80C then transferred to a cryogenic unit 24 hours later until analysis.

### PBMC Phenotyping

Frozen PBMCs were thawed, washed in FACS buffer (2% FBS, 1mM EDTA in PBS), and counted on TC20 (Biorad) before surface staining using two independent flow panels. For the innate panel, the following antibodies were used: CD3 (SP34, BD Pharmingen) and CD20 (2H7, BioLegend) for the exclusion of T & B lymphocytes, respectively. We further stained for CD56 (HCD56, Biolegend), CD57 (HNK-1, BioLegend), KLRG1 (SA231A2, BioLegend) CD16 (3G8, BioLegend), CD14 (M5E2, BioLegend), HLA-DR (L243, BioLegend), CD11c (3.9, ThermoFisher Scientific), CD123 (6H6, BioLegend) and CD86 (IT2.2, BioLegend). For the adaptive panel, the following antibodies were used: CD4 (OKT4, BioLegend), CD8b (2ST8.5H7, Beckman Coulter), CD45RA (HI100, TONBO Biosciences), CCR7 (G043H7, BD Biosciences), CD19 (HIB19, BioLegend), IgD (IA6-2, BioLegend), CD27 (M-T271, BioLegend), KLRG1 (SA231A2, BioLegend) and PD-1 (Eh12.2h7, BioLegend). Cells were stained with Ghost Dye viability dye (TONBO biosciences) for 30 minutes at 4C per manufacturer’s instructions, washed, surface stained with either innate or adaptive panels for 30 minutes at 4C. Samples were then washed and analyzed on Attune NxT Flow Cytometer (ThermoFisher Scientific, Waltham MA).

### Serology

RBD and NP end-point titers were determined using standard ELISA and plates coated with 500 ng/mL SARS-CoV-2 Spike-protein Receptor-Binding Domain (RBD) (GenScript, Piscataway NJ) or 1ug/mL SARS-CoV-2 nucleocapsid protein (NP). Heat inactivated plasma was added in 3-fold dilutions starting at 1:50. Responses were visualized by adding HRP-anti-human IgG (BD Pharmingen) followed by the addition of Phenylenediamine dihydrochloride (ThermoFisher Scientific). ODs were read at 490 nm on a Victor3™ Multilabel plate reader (Perkin Elmer). Batch differences were minimized by normalizing to positive control samples run on each plate.

### Placenta Processing

Placental decidua and villous chorion membranes were separated immediately, immersed in RPMI supplemented with 10% FBS and antibiotics. Samples were processed within 24 hrs of collection. Decidua basalis membranes were washed thoroughly in HBSS to remove contaminating blood. Tissues were minced into approximately 0.2-0.3 mm^3^ cubes and enzymatically digested in 0.5mg/mL collagenase V (Sigma, C-9722) solution in 50 mL R3 media (RPMI 1640 with 3% FBS, 1% Penicillin-Streptomycin, 1% L-glutamine, and 1M HEPES) at 37C for 1 hour. The disaggregated cell suspension was passed through tissue strainers to eliminate large tissue chunks. Cells were pelleted from the filtrate, passed through 70-um cell sieve, centrifuged, and resuspended in R3 media. Red blood cells were lysed using RBC lysis buffer (155 mM NH_4_Cl, 12 mM NaHCO_3_, 0.1 mM EDTA in double-distilled water) and resuspended in 5 mL R3 media. The cell suspension was then layered on a discontinuous 60% and 40% percoll gradient and centrifuged for 30 minutes with brakes off. Immune cells at the interface of 40% and 60% gradients were collected, washed in HBSS, counted, and cryopreserved for future analysis.

### Decidua Immunophenotyping

1-2 × 10^6^ fresh decidual leukocytes were washed with PBS and stained using the following cocktail of antibodies: CD45 (HI30, Biolegend), CD66b (G10F5, BioLegend), CD20 (2H7, BioLegend), CD4 (OKT4, BioLegend), CD8b (2ST8.5H7, Beckman Coulter), CD14 (M5E2, BioLegend), HLA-DR (L243, BioLegend), CD56 (HCD56, Biolegend), CD16 (3G8, BioLegend), CD11c (3.9, ThermoFisher Scientific), CD123 (6H6, BioLegend), for 20 minutes in dark at 4C. Samples were washed twice in FACS buffer and resuspended in 400 uL. All samples were acquired with the Attune NxT Flow Cytometer (ThermoFisher Scientific, Waltham MA), immediately after addition of SYTOX Red Dead Cell Stain (1:1000). Data were analyzed using FlowJO (Ashland OR).

### 3’ multiplexed single cell RNA sequencing

Freshly thawed decidual immune cells (1-2 10e^6^ cells) were incubated with Fc blocker (Human TruStain FcX, Biolegend) in PBS with 1% BSA for 10 minutes at 4C and then surface stained with CD45-FITC (HI30, Biolegend) for 30 minutes in the dark. Samples were then washed twice in PBS with 0.04% BSA and incubated with individual 3’ CellPlex oligos (10X Genomics) per manufacturer’s instructions for 5 minutes at room temperature. Pellets were washed three times in PBS with 1% BSA, resuspended in 300 uL FACS buffer and sorted on BD FACS Aria Fusion into RPMI (supplemented with 30% FBS) following addition of SYTOX Blue stain for live versus dead discrimination. Sorted live cells were counted in triplicates on a TC20 Automated Cell Counter (BioRad), washed and resuspended in PBS with 0.04% BSA in a final concentration of 1500 cells/uL. Single cell suspensions were then immediately loaded on the 10X Genomics Chromium Controller with a loading target of 20,000 cells. Libraries were generated using the V3 chemistry (for gene expression) and Single Cell 3□ Feature Barcode Library Kit per manufacturer’s instructions (10X Genomics, Pleasanton CA). Libraries were sequenced on Illumina NovaSeq with a sequencing target of 30,000 gene expression reads and 5,000 multiplexed CellPlex reads per cell.

### 5’ multiplexed single cell RNA sequencing with feature barcoding

Matched PBMC and decidual leukocytes were thawed, washed, filtered, and stained with Ghost Violet 540 live-dead stain (Tonbo Biosciences) for 30 minutes in the dark at 4C. Samples were washed thoroughly with a cell staining buffer (1X PBS with 0.5% BSA), and Fc blocked for 10 minutes (Human TruStain FcX, Biolegend), and incubated with a cocktail containing CD3-PE (SP34, BD Pharmingen), 0.5 ug each of oligo tagged CD4 (TotalSeq™-C0072, Biolegend), CD8 (TotalSeq™-C0046, Biolegend), CCR7 (TotalSeq™-C0148, Biolegend), CD45RA (TotalSeq™-C0063, Biolegend), CD69 (TotalSeq™-C0146, Biolegend), CD103 (TotalSeq™-C0145, Biolegend), PD-1(TotalSeq™-C0088, Biolegend), CD25 (TotalSeq™-C0085, Biolegend), and unique 5’ hashing antibody (TotalSeq™-C0251, C0254, C0256, and C0260, Biolegend) for an additional 30 minutes at 4C. Samples were washed four times with 1X PBS (serum and azide free), filtered using Flowmi 1000 uL pipette strainers (SP Bel-Art) and resuspended in 300 uL FACS buffer. CD3+ T cells were sorted on the BD FACS Aria Fusion into RPMI (supplemented with 30% FBS). Sorted pellets were counted in triplicates on a TC20 Automated Cell Counter (BioRad), washed and resuspended in PBS with 0.04% BSA in a final concentration of 1500 cells/uL. Single cell suspensions were then immediately loaded on the 10X Genomics Chromium Controller with a loading target of 20,000 cells. Libraries were generated using the 5’ V2 chemistry (for gene expression) and Single Cell 5□ Feature Barcode Library Kit per manufacturer’s instructions (10X Genomics, Pleasanton CA). Libraries were sequenced on Illumina NovaSeq with a sequencing target of 30,000 gene expression reads and 10,000 multiplexed CellPlex reads per cell.

### Single cell RNA-Seq data analysis

For 3’ gene expression with CellPlex, raw reads were aligned and quantified using Cell Ranger (version 6.0.2, 10X Genomics) against the human reference genome (GRCh38) using the *multi* option and CMO information. Only singlets identified from each sample were included in downstream analyses. For 5’ gene expression, alignments were performed using the *feature* and *vdj* option in Cell Ranger. Following alignment, hashing (HTO) and cell surface features (Antibody Capture) from feature barcoding alignments were manually updated in cell ranger generated features file. Doublets were then removed in Seurat using the *HTODemux* function, which assigned sample identity to every cell in the matrix. Droplets with ambient RNA (cells fewer than 400 detected genes), dying cells (cells with more than 20% total mitochondrial gene expression) were excluded during initial QC. Data normalization and variance stabilization were performed on the integrated object using the *NormalizeData* and *ScaleData* functions in Seurat where a regularized negative binomial regression corrected for differential effects of mitochondrial and ribosomal gene expression levels. Dimension reduction was performed using *RunPCA* function to obtain the first 30 principal components and clusters visualized using Seurat’s *RunUMAP* function. Cell types were assigned to individual clusters using *FindMarkers* function with a fold change cutoff of at least 0.4 and using a known catalog of well-characterized scRNA markers for human PBMC and decidual leukocytes. For T cells, 5’ feature barcoding reads were normalized using centered logratio (CLR) transformation. Differential markers between clusters were then detected using *FindMarkers* function. A combination of gene expression and protein markers were used to define T cell subsets. CCR7 staining did not exhibit a significant positive peak, hence was excluded from all downstream analyses. List of cluster specific markers is provided in Supp Table 2. Differential expression analysis was performed using wilcoxon rank-sum test using default settings in Seurat. Only statistically significant genes (log_10_(fold change) cutoff ≥ 0.4; adjusted p-value ≤ 0.05) were included in downstream analyses. Module scores for specific gene sets were incorporated cluster-wise using *AddModuleScores* function.

### scTCR Analysis

TCR reads were aligned to VDJ-GRCh38 ensembl reference using Cell Ranger 6.0 (10X Genomics) generating sequences and annotations such as gene usage, clonotype frequency, and cell specific barcode information. Only cells with one productive alpha and/or one productive beta chain were retained for downstream analyses. CDR3 sequences were required to have length between 5 and 27 amino acids, start with a C, and not contain a stop codon. Clonal assignments from cellranger were used to perform all downstream analysis using the R package immunarch. Data were first parsed through *repLoad* function in immunarch, and clonality examined using *repExplore* function. Family and allele level distributions of TRA and TRB genes were computed using *geneUsage* function. Diversity estimates (Hill numbers) were calculated using *repDiversity* function.

### Statistical analysis

Data sets were first tested for normality using Shapiro-Wilk test. Two way comparisons for normally distributed data were tested for significance using unpaired t-test with welch’s correction. For comparisons involving multiple groups, differences were tested using one-way ANOVA followed by Holm Sidak’s multiple comparisons tests. P-values less than or equal to 0.05 were considered statistically significant. Values between 0.05 and 0.1 are reported as trending patterns.

## RESULTS

### Peripheral and decidual immunological changes in response to asymptomatic/mild SARS-CoV-2 infection

To comprehensively assess the impact of asymptomatic SARS-CoV-2 infection on the immune landscape of the maternal fetal interface, we collected decidua basalis (maternal membrane) and blood at delivery from SARS-CoV-2 PCR+ and/or seropositive women who experience asymptomatic/mild SARS-CoV-2 infection **(Figure 1A and Supp Table 1)**. Blood samples were used to obtain complete blood counts, plasma antibody titers, and assess changes in peripheral immune blood cell (PBMC) composition by flow cytometry **(Figure 1A)**. Decidual immune cells were phenotyped using multispectral flow cytometry and single cell RNA sequencing to assess immune perturbations at the maternal/fetal interface.

**Figure 1:**
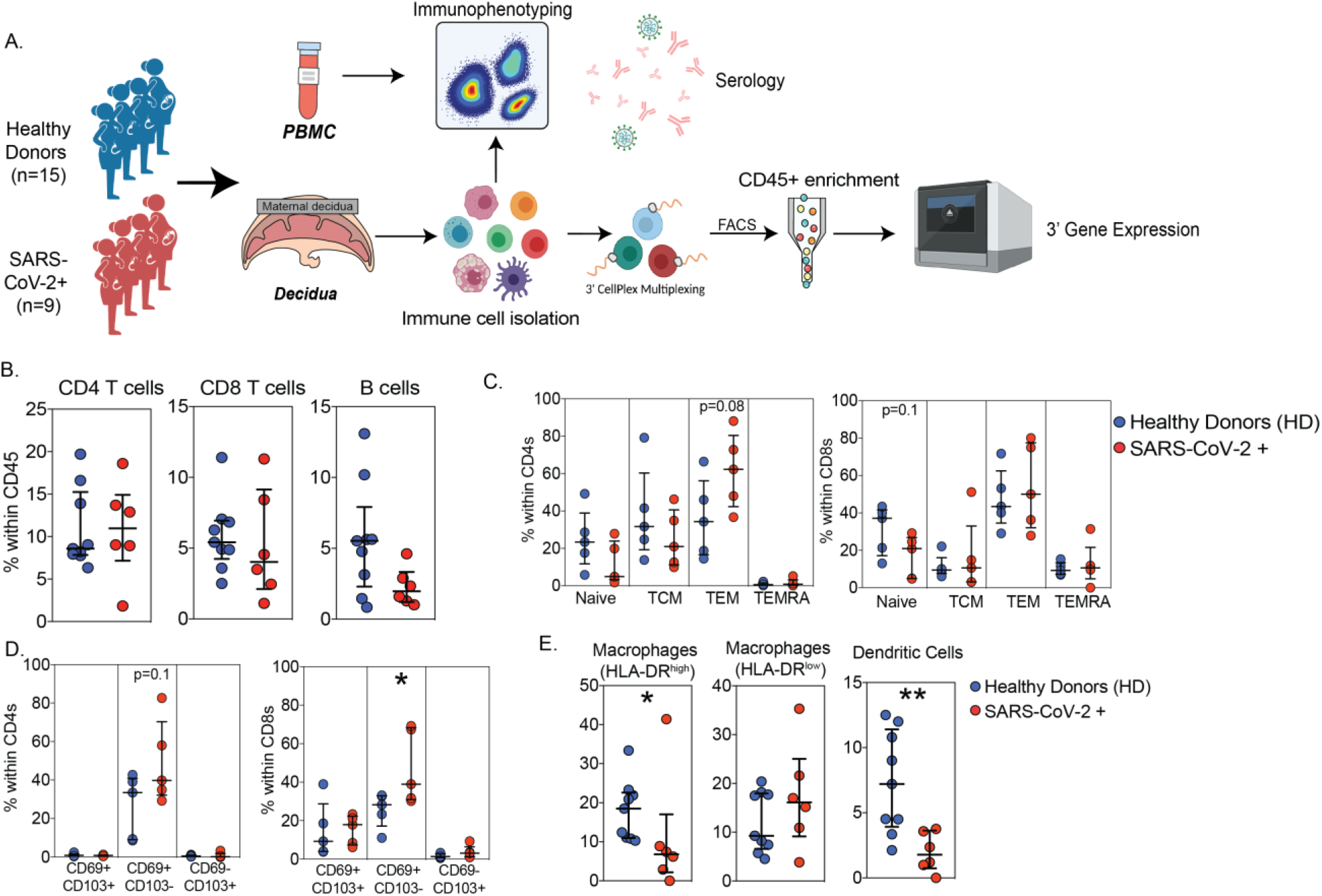
Immunological changes in term maternal decidua with asymptomatic SARS-CoV-2 infection. (A) Experimental design for the study. Placenta and blood were collected from healthy donors (n=15) and SARS-CoV-2 PCR+ or seropositive pregnant women experiencing asymptomatic/mild disease (n=9) at delivery. Immunological changes in PBMC were characterized using multicolor flow cytometry. SARS-CoV-2 specific Endpoint titers were measured in plasma using ELISA. Decidual membranes (decidua basalis) were separated from fetal membranes, processed by mechanical and enzymatic methods to isolate immune cells which were multiplexed and profiled using single cell RNA sequencing. (B-C) Dot plots of decidual (B) T and B cells and (C) T cell subsets (naive, central, and effector memory). (D) Frequencies of tissue resident CD4 and CD8 T cell subsets characterized by CD69 and CD103 expression. (E) Dot plots of frequencies of innate immune cells (macrophages and DCs) within the CD45 compartment of decidua basalis of healthy donors. Two group differences were tested using unpaired t-test with Welch’s correction. Error bars represent median values and interquartile ranges. (p-values: * - p<0.05; ** - p<0.01)

Complete blood cell counts indicated that absolute numbers of circulating granulocytes, monocytes, and platelets increased with infection, but no changes in lymphocyte numbers were detected **(Supp Figure 1A)**. Multicolor flow analysis of PBMC **(Supp Figure 1B)** revealed a redistribution of naïve and memory T cell subsets in the absence of changes in total CD4 and CD8 T cell frequencies **(Supp Figure 1C,D)**. Specifically, infection was associated with reduced abundance of naïve CD4 T cells but modest expansion of central memory (TCM) **(Supp Figure 1D)**. Similar trends were observed in the CD8 subset where a modest reduction in naïve T cells was accompanied by increased frequency of effector (EM) and terminally differentiated effector memory (TEMRA) cells **(Supp Figure 1D)**. Frequency of total B cells and relative abundance of naïve/memory subsets did not vary significantly with infection **(Supp Figure 1E)**. Although frequency of total NK cell proportions did not vary with infection, the relative proportions of CD16^br^CD56^dim^ expanded with infection while those of CD16+CD56+ subset decreased **(Supp Figure 1F)**. Similarly, while frequencies of total monocytes and DC were comparable between the two groups **(Supp Figure 1G)**, a modest expansion of CD16++ non-classical monocytes and a significant drop in plasmacytoid DC (pDC) was noted **(Supp Figure 1H)**. Finally no differences in surface expression of MHC-Class II molecule HLA-DR **(Supp Figure 1I)** or activation marker CD86 **(Supp Figure 1J)** were observed with infection.

Flow analysis of decidual leukocytes revealed no changes in total T or B cell frequencies **(Figure 1B and Supp Figure 2A)**. However, we observed a modest expansion of CD4 effector memory cells and modest reduction in the CD8 naive T cells **(Figure 1C)** as well as frequencies of CD69+CD103-CD4 and CD8 tissue resident T cells with asymptomatic/mild disease **(Figure 1D)**. No differences in total decidual NK cell frequencies or subsets were seen with SARS-CoV-2 infection **(Supp Figure 2B,C)**. However, proportions of both HLA-DR^high^ macrophages and DCs decreased significantly with asymptomatic infection **(Figure 1E)**.

**Figure 2:**
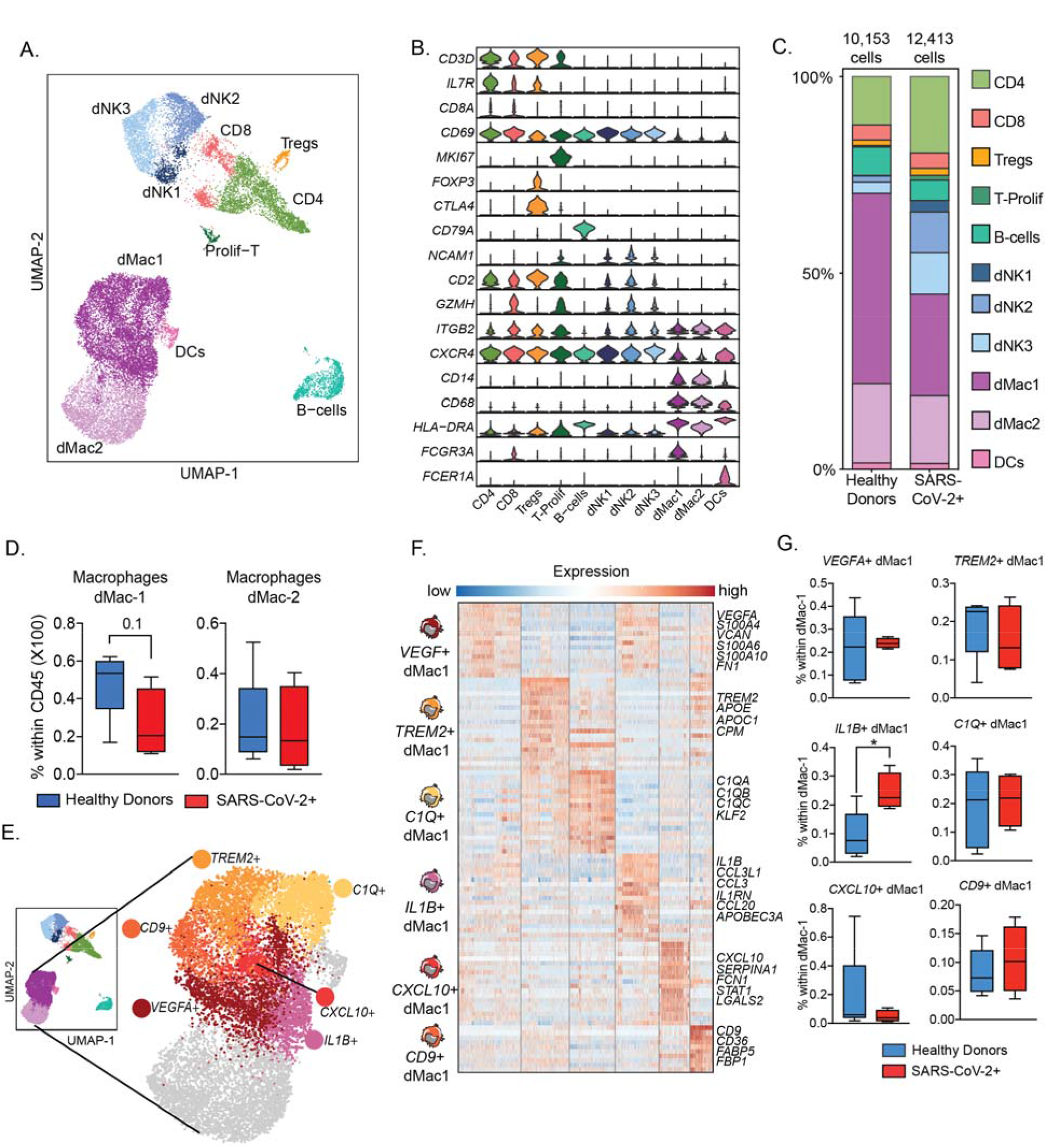
Unsupervised analysis of CD45+ compartment within term maternal decidua and adaptations with asymptomatic SARS-CoV-2 infection. (A) Uniform Manifold Approximation and Projection (UMAP) representation of 22,566 immune cells within the term decidual CD45+ compartment showing 11 clusters. (B) Violin plots of key gene markers used for cluster annotations. Y axis represents the normalized transcript counts. (C) Stacked bar graph comparing distribution of decidual immune cell clusters in healthy donors (10,153 cells) and mothers with asymptomatic/mild SARS-CoV-2 infection (12,413 cells). (D) Box and whiskers comparing single cell frequencies of two macrophage subsets dMac1 and dMac2 with asymptomatic SARS-CoV-2 infection. (E) UMAP of term decidual myeloid cells highlighting the diverse states within dMac1 (HLA-DRAhigh) macrophage subset. Each color indicates one of 6 dMac1 clusters identified with a prominent marker highlighted. (F) Heatmap of normalized transcript counts of top 20 genes within each of the 6 dMac1 clusters. A handful of highly expressed markers within each cluster are highlighted. Color in the heatmap represents scaled average expression ranging from low (blue) to high (red). Box and whiskers comparing changes in macrophage subset frequencies with asymptomatic SARS-CoV-2 infection. Two group differences were tested using unpaired t-test with Welch’s correction. (p-values: * - p<0.05)

### Unsupervised single cell analysis of decidual leukocytes reveals heightened myeloid cell activation with asymptomatic SARS-CoV-2 infection

We next assessed rewiring of immune (CD45+) cell states with asymptomatic infection in an unbiased way at the single cell resolution. Decidual leukocytes from 4 individuals per group multiplexed using lipid tagged oligos (compatible with 3’ single cell gene expression), sorted to remove dead cells, and analyzed using single cell RNA sequencing (scRNA seq) **(Figure 1A)**. Following doublets and dead cell removal, dimension reduction and clustering, our analysis identified 11 unique immune cell clusters **(Figure 2A)**. Cluster annotations were derived from differential marker analysis **(Figure 2B)** and confirmed using markers previously described for first trimester decidual immune landscape (16). Within the lymphoid clusters, B cells were identified based on high expression of *CD79A*; CD4 T based on *IL7R*; CD8 T on *CD8A* and *GZMH* expression; regulatory T cells on *FOXP3* and *CTLA4* expression; proliferating T cells on CD2, CD3 and *MKI67* expression; and NK cell subsets on differential expression of *CSF1, GZMH, CXCR4*, and *ITGB2* (16) **(Figures 2A and 2B)**. Myeloid clusters consisted primarily of macrophages, which were broadly divided by magnitude of *HLA-DRA* expression (as observed in flow cytometry, **Supp Figure 2A**) into dMac1 (*HLA-DR*^high^) and dMac2 (*HLA-DR*^low^) (16) and DCs, expressing high *FCER1A* **(Figures 2A and 2B)**.

We next stratified clusters by infection status **(Figure 2C and Supp Figure 3A)** and patient of origin (since the cells from each subject were barcoded using oligo tagged lipids), and compared cluster frequencies between the two groups **(Supp Figure 3B)**. No changes were observed in the relative frequency of decidual T cells **(Supp Figure 3C, 3D)**, B cells **(Supp Figure 3E)**, or NK cells **(Supp Figure 3F)**. Additional analysis shows a modest increase in dNK2 (16) **(Supp Figure 3G)**; therefore, we identified genes differentially expressed with SARS-CoV-2 infection within decidual NK cells. Modest gene expression changes (73 up and 33 down with infection) were identified with pathways associated with protein folding, MAPK signaling, and type-I interferon signaling up-regulated with infection, whereas immune activation, adhesion associated terms were down-regulated **(Supp Figure 3H)**

**Figure 3:**
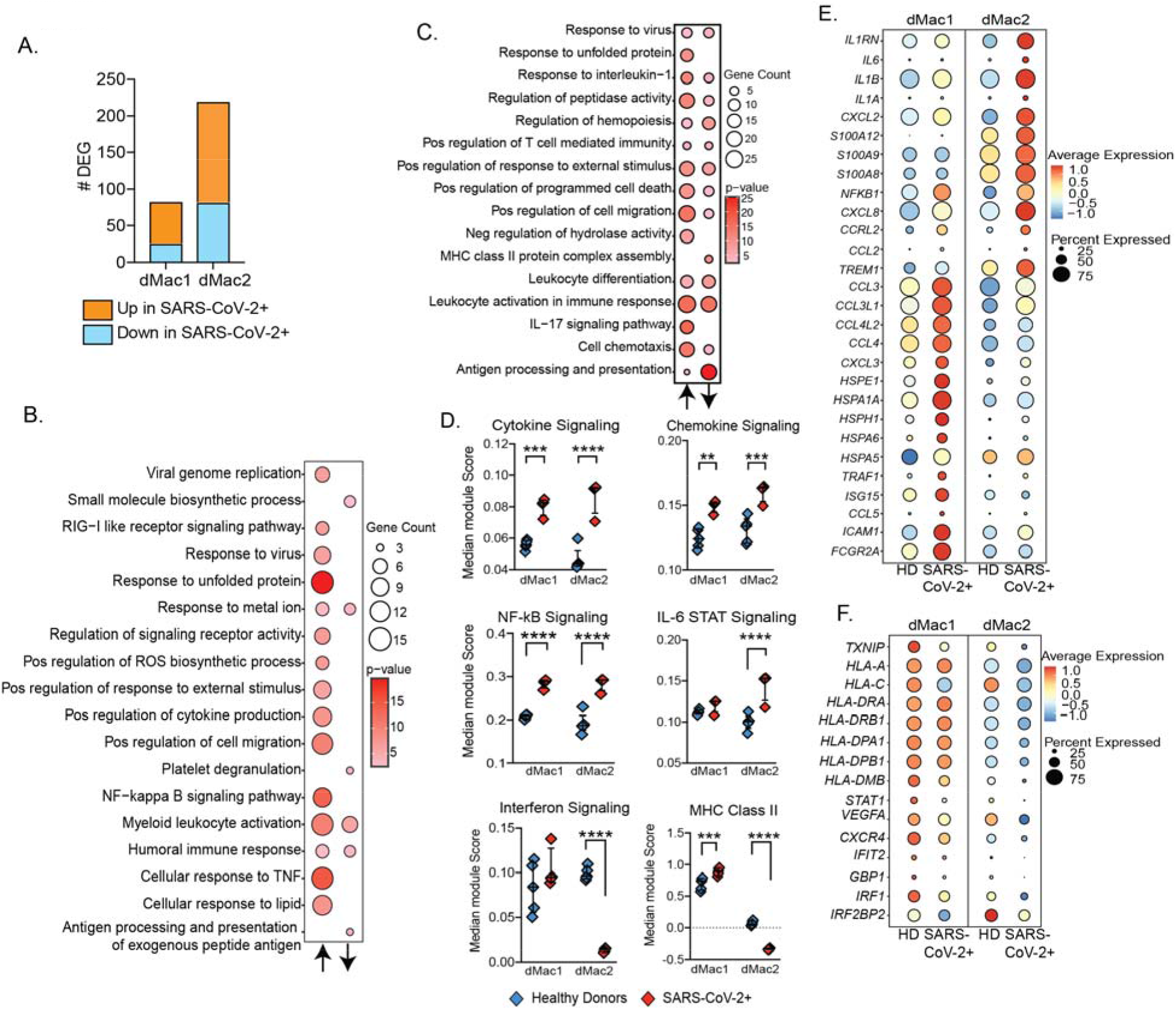
Asymptomatic SARS-CoV-2 infection and associated myeloid cell activation in term decidua. (A) Differentially expressed genes in decidual macrophage subsets with asymptomatic SARS-CoV-2 infection. (B-C) Bubble plots representing two-way functional enrichment of genes up- and down-regulated with asymptomatic/mild SARS-CoV-2 infection in (B) dMac1 and (C) dMac2 macrophage subsets. Size of the bubble represents the number of genes mapping to the term whereas color represents level of statistical significance. Two-way enrichments were performed in Metascape. (D) Dot plots comparing changes in module scores of innate immune pathways in decidual macrophage subsets with asymptomatic/mild SARS-CoV-2 infection. Each dot represents the median module score for the individual. Error bars represent median values and interquartile ranges. (E-F) Bubble plot depicting normalized expression of key differentially expressed genes (statistically significant adjusted p<0.05 and fold change>0.4). (E) upregulated and (F) downregulated with infection. Size of the bubble represents percentage of cells within the cluster (dMac1 or dMac2) expressing the gene, while color represents the average expression (normalized transcripts) scaled from low (blue) to intermediate (yellow) to high (red). (p-values: ** - p<0.01; *** - p<0.001; **** - p<0.0001)

Interestingly, and in contrast to the flow cytometry data presented above, no differences in DC frequencies were detected from the scRNA Seq data **(Supp Figure 3I)**. However, as observed with flow cytometry, we observed a selective loss of *HLA-DR*^high^ macrophages (dMac1) with asymptomatic SARS-CoV-2 infection **(Figure 2D)**. Additional investigation on this subset revealed a high degree of heterogeneity, with the presence of 6 distinct macrophage states **(Figure 2E)**. These included regulatory macrophages expressing *TREM2*, and activated subsets expression high levels of complement genes (*C1Q*+), tetraspanin *CD9*, pro-inflammatory cytokine *IL1B*, chemokine *CXCL10* (IP-10), or growth factor *VEGFA* **(Figures 2E, F)**.

Although frequencies of dMac1 were reduced with infection **(Figure 2G)**, *IL1B*+ dMac1 subset expressing cytokines *IL1B, CCL3, CCL20* expanded with infection **(Figure 3H)**. Moreover, robust gene expression changes were observed within both macrophage subsets in response to infection **(Figure 4A)**. In dMac1 macrophages, infection was associated with the induction of genes involved in the antiviral response, viral sensing, NF-*κ*B signaling, and cytokine production, while a small number of genes that play a role in myeloid cell activation and antigen processing and presentation were suppressed **(Figure 3B)**. More robust gene expression differences were observed in dMac2 subset **(Figure 3A)**, altering signatures associated with immune activation, upregulating genes involved in chemotaxis, cell death, and IL-17 signaling, and suppressing genes involved in antigen processing and presentation **(Figure 3B)**.

**Figure 4:**
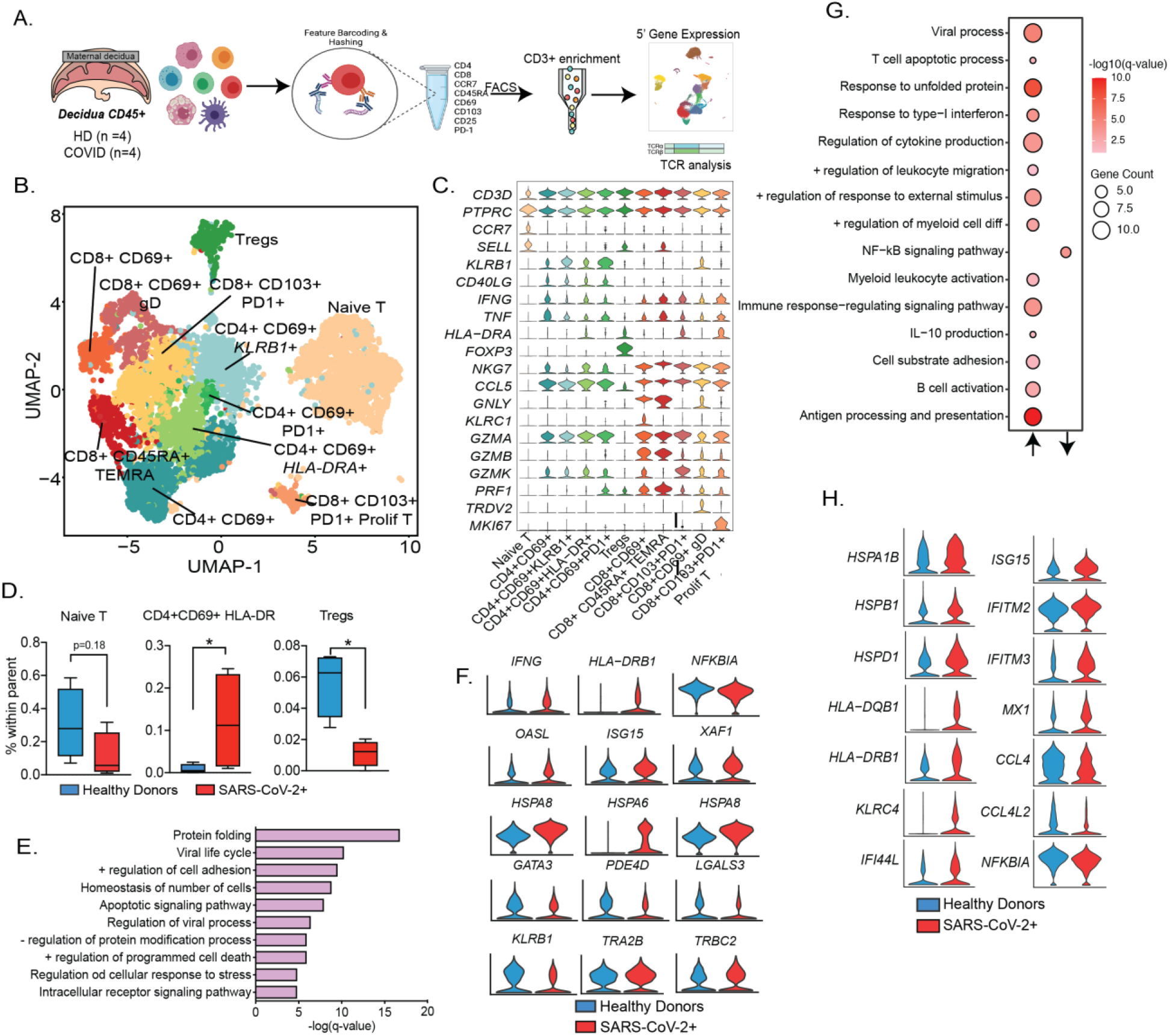
Decidual T cell adaptations to maternal asymptomatic SARS-CoV-2 infection. (A) Experimental design for deep profiling of T cells from blood and maternal decidua. Isolated immune cells from decidua basalis were surface stained with fluorescent marker for T cells (CD3), DNA oligo tagged antibodies for CD4, CD8, naive, and memory, and tissue resident markers, along with hashing antibodies, enriched for T cells using FACS and subjected to 5’ gene expression and TCR analysis (immune repertoire) on the 10X platform. (B) Dimension reduction and clustering of decidual T cells identifies 11 distinct T cell clusters at term, annotated by unique surface and gene expression markers. (C) Stacked violin plots highlighting key gene expression markers of T cell subsets within term maternal decidua. Y axis represents the normalized transcript counts. (D) Box and whiskers plot of decidual T cell subsets changing with asymptomatic SARS-CoV-2 infection. (E) Functional enrichment of top genes (log2 fold change ≥ 0.4, q-value ≤ 0.05) within the CD4+CD69+HLA-DRAhigh T cell cluster. Y axis represents q-values (negative log 10 transformed). (F) Violin plots of select statistically significant (log2 fold change ≥ 0.4, up or downregulated, q-value ≤ 0.05) differentially expressed genes with asymptomatic SARS-CoV-2 infection within memory CD4 T cell clusters (G) Bubble plots representing two-way functional enrichment of genes up- and down-regulated with asymptomatic SARS-CoV-2 infection in memory CD8 T cell clusters. Size of the bubble represents the number of genes mapping to the term whereas color represents level of statistical significance. (H) Violin plots of select statistically significant (log2 fold change ≥ 0.4, up or downregulated, q-value ≤ 0.05) differentially expressed genes with asymptomatic SARS-CoV-2 infection within memory CD4 T cell clusters. Two group differences were tested using unpaired t-test with Welch’s correction. (p-values: ** - p<0.01; *** - p<0.001; **** - p<0.0001)

We next compared scores for specific signaling pathways across both groups. SARS-CoV-2 infection was associated with an increase in module scores of cytokine and chemokine signaling **(Figure 3C)**, notably NF-*κ*B, TNF and IL-17 signaling **(Supp Figure 4A)** and pathogen sensing pathways such as TLR and RIG-I signaling **(Supp Figure 4B)**. The higher chemokine module score is illustrated by increased expression of several chemokine genes (such as *CCL3, CCL3L1, CCL4, CXCL3*, and *CCL5*), notably in dMac1 subset **(Figure 3E)**. Interestingly, only dMac2 subset exhibited heightened IL-6 STAT signaling **(Figure 3D)** including an up-regulation of inflammatory genes such as alarmins (*S100A8, S100A9, S10012*), *TREM1*, and cytokines (*IL1A, IL1B, IL6*) with infection **(Figure 3E)**. On the other hand, only dMac2 macrophage subset exhibited dampened type-I interferon signaling **(Figure 3D)** with key interferon stimulated genes (*IFIT2, GBP1, IRF1*) down regulated with infection **(Figure 3F)**.

Finally, we observed a differential impact of mild infection of MHC signaling in decidual macrophage subsets. Module scores for MHC class II signaling was down-regulated in dMac2 macrophages (*HLA-DRA, HLA-DRB1, HLA-DPB1*) **(Figures 3D, 3F)**, while MHC class I signaling was down-regulated in both subsets **(Figure 3F, Supp Figure 4C)**. Interestingly, surface expression of activation markers CD40 and CD86 did not change with infection **(Supp Figure 4D)**. On the other hand, a higher expression level of M2 marker CD206 was detected on dMac1 macrophages, and modest increases in interferon receptor IFNAR1 expression was seen on both subsets **(Supp Figure 4E)**.

### Expansion of activated CD4 T cells and upregulation of type-I interferon signaling in CD4 and CD8 T cells following asymptomatic SARS-CoV-2 infection

To investigate the impact of asymptomatic SARS-CoV-2 infection in pregnant mothers on diverse T cell states within term placentas, we profiled CD3+ sorted cells from decidual leukocytes and performed single cell analysis (gene expression and T cell repertoire) following multiplexing and feature barcoding, staining for CD4, CD8, CD45RA, tissue markers CD69 and CD103, and regulatory markers PD1 and CD25. Orthogonal readouts were measured in T cells from matched blood samples **(Figure 4A)**. After doublets and ambient RNA removal, we identified 10 memory T cell clusters in addition to a naive cluster **(Figure 4B)** within the decidua. Both surface expression of CD4, CD8, CD69, CD103, CD25, and PD-1 from feature barcoding **(Supp Figure 5A)** and highly expressed RNA markers **(Figure 4C)**, were used to annotate these memory T cell clusters. All memory CD4s expressed CD69, with three additional clusters expressing either high levels of PD-1, or RNA levels of *HLA-DRA* or *KLRB1*. Memory CD8 T cells on the other hand expressed either CD69 (including a subset of gamma delta T cells) or CD103 and PD-1 (including a subset of proliferating CD8s) **(Figures 4B, 4C, and Supp Figure 5A)**.

**Figure 5:**
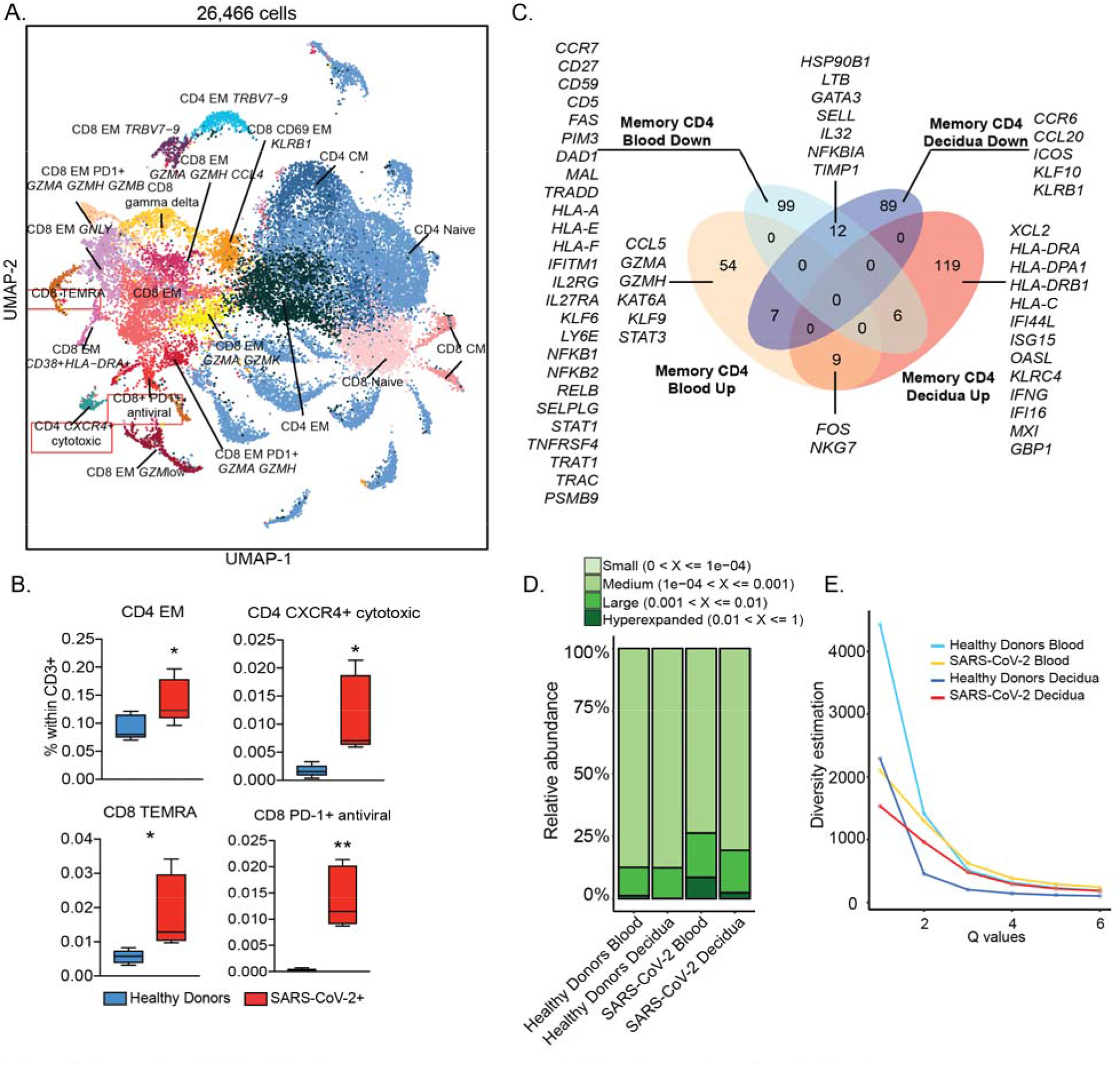
Comparing decidual T cell responses to asymptomatic infection to adaptations in blood. (A) Dimension reduced UMAP representation of 26,466 blood T cells collected during delivery – 12,819 from healthy donors and 13,647 from mothers with asymptomatic SARS-CoV-2 infection. Key clusters are annotated based on top gene expression and protein markers identified by Seurat. (B) Box and whiskers plot of key blood CD4 (top) and CD8 (bottom) T cell subsets changing with asymptomatic SARS-CoV-2 infection. (C) Venn diagram comparing differentially expressed genes (log2 fold change ≥ 0.4, up or downregulated, q-value ≤ 0.05) in memory CD4 T cells from blood and decidua. Select genes within this comparison are highlighted. (D) Stacked bars comparing T cell clone sizes (based on proportions within clonal T cells) in blood and decidua in healthy donors and mothers with asymptomatic infection. (E) Diversity profiles of T cells with infection – y-axis represents Hill diversity, interpreted as the effective number of clonotypes within the dataset. (p-values: * - p<0.05)

We next compared the proportions of decidual T cell clusters with infection **(Supp Figures 5B and 5C)**. Modest reduction in naive T cells and a statistically significant reduction in regulatory T cell frequencies were observed with infection **(Figure 4D and Supp Figure 5C)**. This reduction was accompanied by an overall expansion of CD69+ memory subsets **(Supp Figures 5C and 5D)**, as observed with flow cytometry **(Figure 1D)**. Interestingly, the subset of activated CD4+CD69+*HLA-DRA*+ significantly expanded with infection **(Figure 4D)**. This subset of T cells, which was almost absent in healthy donors, expressed high levels of genes involved in regulation of viral processes, stress response, cell adhesion, and apoptosis **(Figure 4E)**. Differential gene expression analysis of memory CD4 T cells showed upregulation of *IFNG*, antiviral genes (*OASL, ISG155, XAF1*), heat shock proteins (*HSPA6, HSPA8*), but downregulation of Th2 genes (*GATA3, PDE4D*) **(Figure 4F)**. Although abundance of CD8 T cells subsets did not vary significantly with infection **(Supp Figure 5E)**, differential gene expression analysis revealed elevated signatures of T cell apoptosis, viral processes, myeloid leukocyte activation, and antigen processing and presentation **(Figures 4G and 4H)**. On the other hand, genes associated with NF-*κ*B signaling were down-regulated with infection (*NFKBIA, CCL4*) **(Figures 4G and 4H)**. Finally, increased module scores for IL-2, type I interferon and T cell signaling as well as T cell activation and exhaustion were observed with SARS-CoV-2 infection **(Supp Figure 5F)**.

### Distinct impact of asymptomatic SARS-CoV-2 infection during pregnancy on peripheral and decidual T cells

We next investigated peripheral T cell adaptations to asymptomatic SARS-CoV-2 infection and compared it to changes in placental T cells. Dimension reduction and clustering identified naïve, central memory, and several clusters of effector memory T cells, particularly within CD8 EM **(Figure 5A)** that were annotated using a combination of gene expression **(Supp Figures 6A)** and protein markers **(Supp Figure 6B)**. Several clusters of naïve CD4 T cells varying in their TCR gene usage (*TRA* or *TRB*) were all pooled as naïve CD4 cluster **(Figure 5A)**. We compared frequencies of T cell clusters with infection and carried differential gene expression changes within total memory T cells. Within CD4 T cells, this analysis revealed an expansion of effector memory cells and a cluster of CXCR4+ cells expressing cytotoxic molecules with infection **(Figure 5B)**. Differentially expressed genes with infection in the blood were quite distinct from ones observed in decidua **(Figure 5C)**. For example, while decidual T cells exhibited upregulation of antiviral (*IFI44L, ISG15, OASL, IFNG, MX1, GBP1*) and MHC class II signaling (*HLA-DRA, HLA-DRB1*), matched blood T cells exhibited elevated cytotoxic signatures (*CCL5, GZMA, GZMH*). More importantly, genes associated with apoptotic (*FAS, PIM3, DAD1, MAL, TRADD*), TNF (*CD27, TNFRSF4, STAT1*), and TCR signaling (*TRAT1, TRAC, PSMB9*) were exclusively downregulated in blood CD4 T cells **(Figure 5C)**.

Within CD8+ T cells, infection was associated with an expansion of TEMRA and PD1+ CD8 T cells expressing antiviral transcripts **(Figure 5B, Supp Figure 6C)**. Additionally, distinct antiviral signatures were upregulated in decidua (*ISG15, MX1, IFI6, IFITM2*) compared to blood (*OASL, CXCR4, IFIT2, TNF, TNFAIP3, FOS*) **(Supp Figure 6D)**. Infection was associated with up regulation of MHC class II genes in both decidual and blood CD8 T cells **(Supp Figure 6D)**. However, blood CD8 T cells exhibited profound dampening of cytotoxicity genes such as *GNLY, NKG7, PRF1, KLRD1*, and *KLRK1*. Furthermore several genes involved in TCR signaling (*CD247, LCK, TRAC, TRAB1, CSK, UBB*) were significantly downregulated in blood. Finally, clonal analysis of T cells revealed expansion of large clones (>100 cells) with infection in blood but not in decidua **(Figure 5D and Supp Figure 6E)**, where only small clones were detected. Furthermore, T cells repertoire diversity in blood was lower than that of decidual T cells after infection **(Figure 5E)**.

## DISCUSSION

Placental immune cells facilitate implantation, fetal growth, tolerance, and promote labor (16). Immune responses at the maternal-fetal interface are highly fine-tuned, balancing protecting the fetus from pathogens while also limiting excessive inflammatory exposure associated with stress or infection (17). Recent studies have suggested evidence of immune activation at the maternal fetal interface with severe COVID-19 in the absence of active infection or viral RNA in the placenta (20). Interestingly, a study involving 991 participants concluded that a vast majority of pregnant women with COVID-19 develop asymptomatic or mild version of the disease, with a more prolonged course of disease compared to infection in non-pregnant women (25). However, it is currently unclear if the robust immunological changes at the maternal-fetal interface observed with severe disease exist with mild/asymptomatic disease, and if these changes are linked to peripheral immune response to infection. Therefore, in this study, we collected blood and placentas from pregnant mothers who experienced asymptomatic/mild SARS-CoV-2 infection during pregnancy or were PCR+ at delivery. We interrogated immunological changes in maternal decidua and PBMC using flow cytometry and single cell RNA sequencing.

Mild disease in nongravid population is characterized by lymphopenia, albeit with a lower magnitude than that observed with severe disease (26). Patients with moderate disease also experience an early increase in cytokines, but a progressive reduction in type-I and type-III interferon responses in (27). However, compared to patients with severe COVID-19, circulating proteome of patients who recovered from mild disease enriched for several reparative growth factors (27). Furthermore, while innate immune cell frequencies (monocytes, neutrophils, eosinophils) increase with SARS-CoV-2 infection, this increase was less dramatic with moderate disease, suggesting a less inflammatory immune response with mild/moderate disease (27).

In line with these previous observations, asymptomatic/mild disease during pregnancy was associated with elevated granulocyte, monocytes, and platelet frequencies but with no overt signs of lymphopenia. As reported in most COVID-19 studies (28, 29), mild disease was associated with a reduction in plasmacytoid DCs (pDCs). However, the overall innate immune response to infection appears to be highly attenuated with infection during pregnancy. First, we did not observe a redistribution of monocyte subsets, with the exception of a modest increase in non-classical (CD16++) monocytes. Myeloid DC frequency and their surface expression of HLA-DR and CD86 remained unchanged with disease. These findings are in line with a recently profiled blood immune cells and plasma in pregnant women with asymptomatic disease, where limited immune changes in PBMC composition and cytokine milieu were observed (13). We posit that these differences are driven by Th2 bias observed with pregnancy that may limit inflammation and reinforce immune tolerance (11). Indeed, in support of this hypothesis plasma of mothers with asymptomatic disease had elevated levels of regulatory IL-1RA and IL-10 (13). This immune adaptation with pregnancy does not preclude pregnant women from developing an effective adaptive immune response to the virus since T cells from pregnant mothers with asymptomatic disease have been shown to be functionally competent with respect to *ex vivo* cytokine production and proliferation (13). Our data indicate reduction in naïve CD4 T cells, and expansion of CD8 memory T cells (TEMRA), in addition to enrichment of cytotoxic CD4 and PD1+ antiviral CD8 T cells with infection. Furthermore, serological studies confirmed presence of both RBD and NP end point titers in the enclosed study participants. Whether pregnancy alters the magnitude and/or quality of antigen specific B and T cell responses to SARS-CoV-2 will need to be the focus of future research.

Severe COVID-19 disease during pregnancy has been recently shown to be associated with myeloid and T cell activation in the placenta (20). However studies on mild disease are lacking. While we report broad myeloid cell activation in both macrophage subsets with asymptomatic disease, our analyses revealed differential outcomes within the two decidual macrophage subsets– HLA-DR^high^ (dMac1, tissue resident decidual macrophages) and HLA-DR^low^ (dMac2, blood monocyte derived decidual macrophages) (16). First, asymptomatic/mild disease resulted in selective loss of the tissue resident dMac1. However, within this cluster of diverse cell states, we observed an increase in cytokine producing (*IL1B*+) macrophages. Given the absence of infection in the placenta, this enrichment of cytokine producing macrophages is most likely a local response to decidual T cell activation or systemic inflammatory cues. Secondly, both macrophage subsets exhibited elevated signatures of TLR, NF-*κ*B, cytokine, and chemokine signaling, suggesting a heightened activation state. However, expression of several heat shock proteins (*HSPA5, HSPA6, HSPE1*) were only upregulated in dMac1, a potentially protective mechanism to cope with cellular stress at the maternal fetal interface. Thirdly, while viral sensing pathways (RIG-I) signaling and interferon receptor expression (IFNAR1) were upregulated with mild/asymptomatic disease, interferon signaling module scores were attenuated in monocyte derived macrophages (dMac2) in the decidua. This observation is in line with several studies on peripheral monocytes from mild/moderate non-gravid subjects, where lower plasma IFNα and lower IFN signaling has been linked to attenuated TNFα and IFNγ in plasma and ensuing cytokine storm. Finally, reduction in HLA-DR expression is a consistent observation of several COVID-19 studies (30, 31), both in mild and severe nongravid subjects. Our analysis suggests significant downregulation of MHC class II signatures in dMac2 but not dMac1 suggesting differential burden of infection on blood derived vs. tissue resident myeloid cells. Monocytes in blood undergo constant turnover, and given the lack of changes in blood monocytes, we argue that SARS-CoV-2 infection results in sustained changes in placental macrophages. These data also suggest that the kinetics by which inflammation is resolved in blood and tissue myeloid compartments are to a large extent, decoupled. Macrophages in the decidua play diverse roles ranging from clearance of apoptotic bodies, wound healing to host defense, pathogen clearance, and facilitation of labor cascade. Thus, an aberrantly activated macrophage state can affect trophoblast function and placental development, contributing to the pathophysiology of fetal growth restriction, pre-eclampsia, preterm birth, and even fetal loss (32-34).

The dichotomy of immune responses in blood versus decidua is more evident in T cells. While T cells in pregnant mothers with asymptomatic COVID-19 have been shown to be functional (13), activated CD4 and CD8 T cells were detected in both blood and decidua. Modest loss of naïve T cells and expansion of memory subsets was observed in both blood and decidua. A recent study in hospitalized patients with severe COVID-19 reported increased expression of *CD69* in decidual T cells (20). Our analysis also expansion of CD8+CD69+ and CD4+CD69+ tissue resident T cells although our participants experience asymptomatic/mild disease. With asymptomatic disease, we observed loss of regulatory T cells in decidua but expansion of HLA-DRA expressing activated CD4 T cells, suggesting a tipping of balance towards Th1 responses. The impact of infection on blood T cells was far more dramatic, with enrichment of effector memory T cells (36, 37) and antiviral PD-1+ CD8 T cells, previously described in nongravid patients (38). Additionally, upregulation of MHC class II molecules on CD8 T cells was reported exclusively in the decidua, whereas a the appearance of cytotoxic CD4 T cells expressing *CXCR4* and high levels of *GZMA, GZMH, CCL5* was seen only in the blood. These observations are consistent with the fact that T cells in the decidua are more poised towards Th2, Th9, Th17 and Tregs compared to circulating T cells (39). Blood helper T cells however, had downregulated signatures of apoptotic signaling, TNF and IFNγ signaling, not observed in decidua. Finally, T cell repertoire analysis, suggests that unlike in blood, T cells in decidua undergo minimal clonal expansion following asymptomatic infection. Taken together, these findings suggest that while antiviral cytotoxic responses are likely restricted to the blood, activated tissue resident decidual T cells are expanded with infection and exhibit signs of heightened interferon signaling.

We acknowledge the limitations of our study that include a small sample size, which combined with the wide window of infectivity during pregnancy, precluded the assessment of gestational age on infection outcomes. Given the limited amount of blood obtained, we were unable to conduct a comprehensive measurement of antigen specific T and B cell responses. Finally, it remains unclear if the placenta harbors virus-specific T cells that migrate from blood. Importantly, while there is no evidence of vertical transmission (24), enrichment of activated T cells and loss of regulatory tissue resident macrophages (dMac1) and Tregs with infection skews the balance of decidual immune cells towards a proinflammatory state. The resulting aberrant immune activation in the placenta and misfiring of local cytokine networks can potentially contribute to pregnancy complications. Additionally, the developing fetal immune system is acutely sensitive to inflammation and stress, and it remains to be seen if such exposures, even with mild maternal infection can have long-term consequences on immunity in the offspring.

---

**Supp Figure 1:**
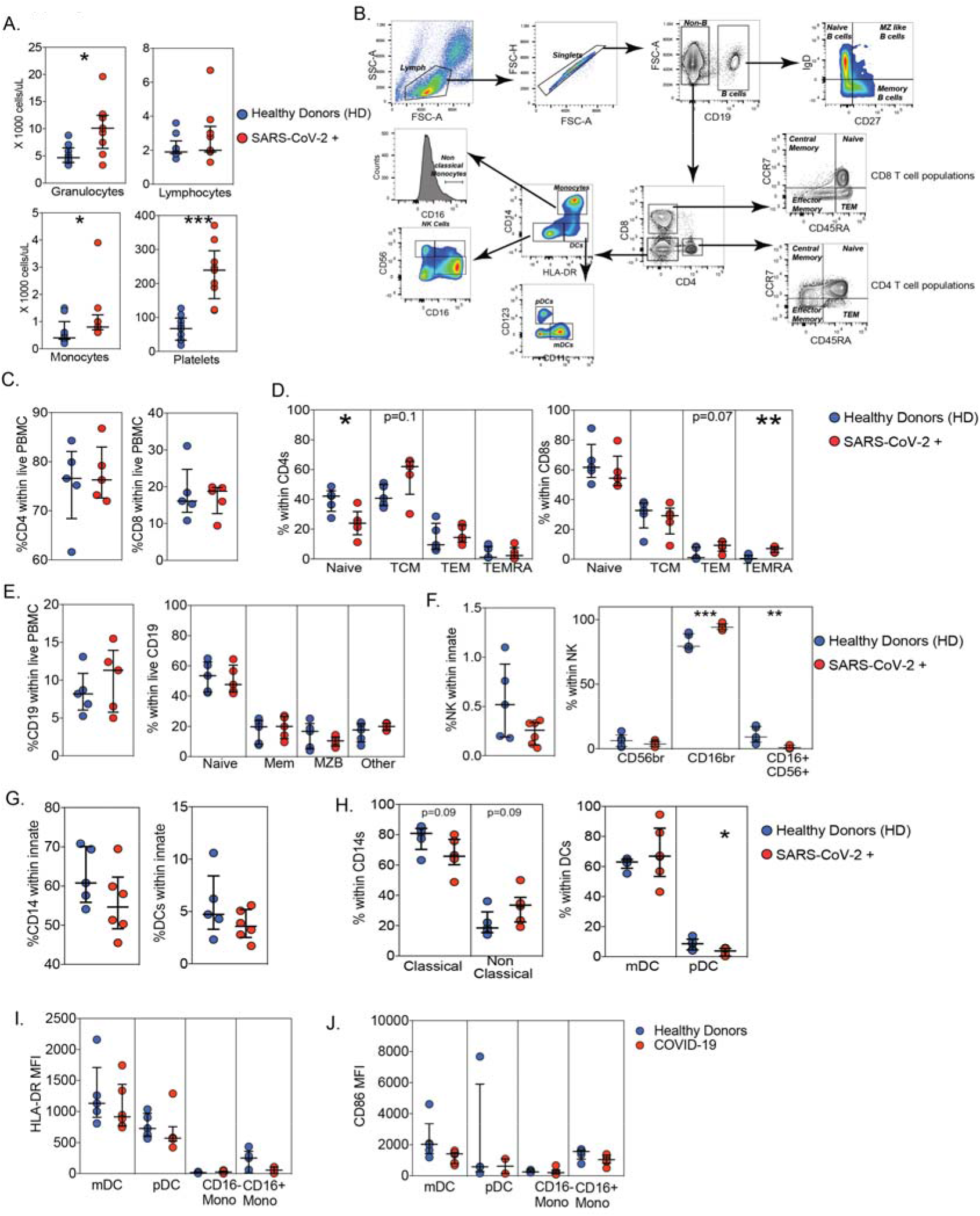
Immunological changes in blood with asymptomatic SARS-CoV-2 infection. (A) Complete Blood Counts (CBC) of samples collected at delivery from pregnant healthy donors and those with a positive PCR or seropositive for SARS-CoV-2 experiencing an asymptomatic/mild disease. (B) Gating strategy for phenotyping peripheral blood mononuclear cells (PBMC) in blood collected at delivery from healthy donors and mothers with asymptomatic SARS-CoV-2 infection. (C-D) Dot plots comparing (C) blood CD4 and CD8 within PBMC and (D) T cell subsets within CD4 and CD8 compartments with maternal asymptomatic infection. (E-F) Frequencies of blood (E) B cells and (F) NK cells and their subsets with asymptomatic infection (n=5/group). (G-H) Dot plots comparing (G) blood monocytes and dendritic cells proportions and (H) their subsets with maternal asymptomatic infection (n=5/group). (I-J) Median fluorescence intensities (MFI) of (I) HLA-DR and (J) CD86 within monocyte and DC subsets. Two group differences were tested using unpaired t-test with Welch’s correction. Error bars represent median values and interquartile ranges. (p-values: ** - p<0.01; *** - p<0.001; **** - p<0.0001)

**Supp Figure 2:**
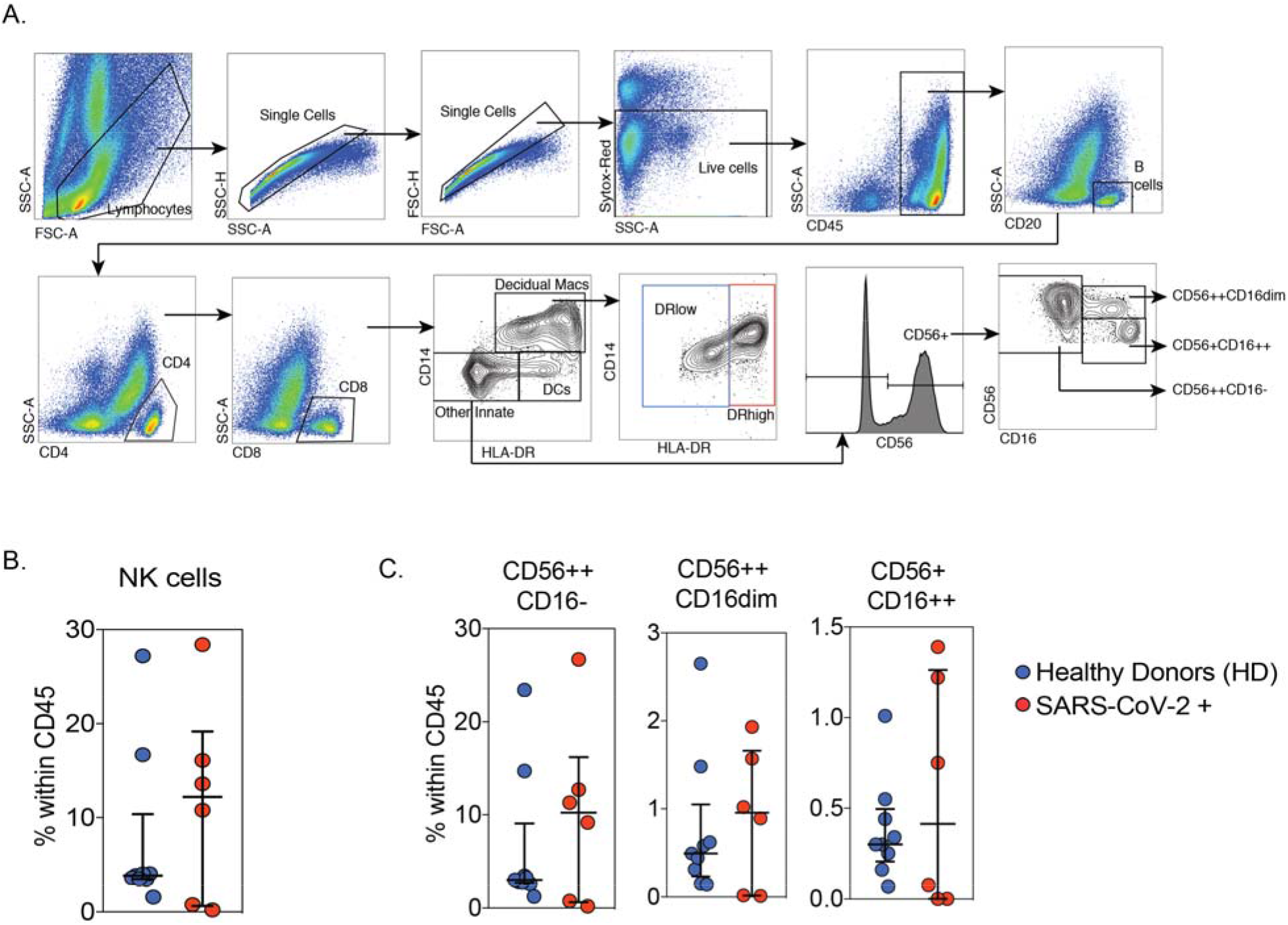
Decidual immune adaptations with asymptomatic SARS-CoV-2 infection during pregnancy. (A) Gating strategy for phenotyping immune cells (CD45+) within term maternal decidua using flow cytometry. (B-C) Dot plots of three decidual NK cell subsets within the CD45+ leukocytes obtained from healthy donors and those with a positive PCR or seropositive for SARS-CoV-2 experiencing an asymptomatic/mild disease. Error bars represent median values and interquartile ranges.

**Supp Figure 3:**
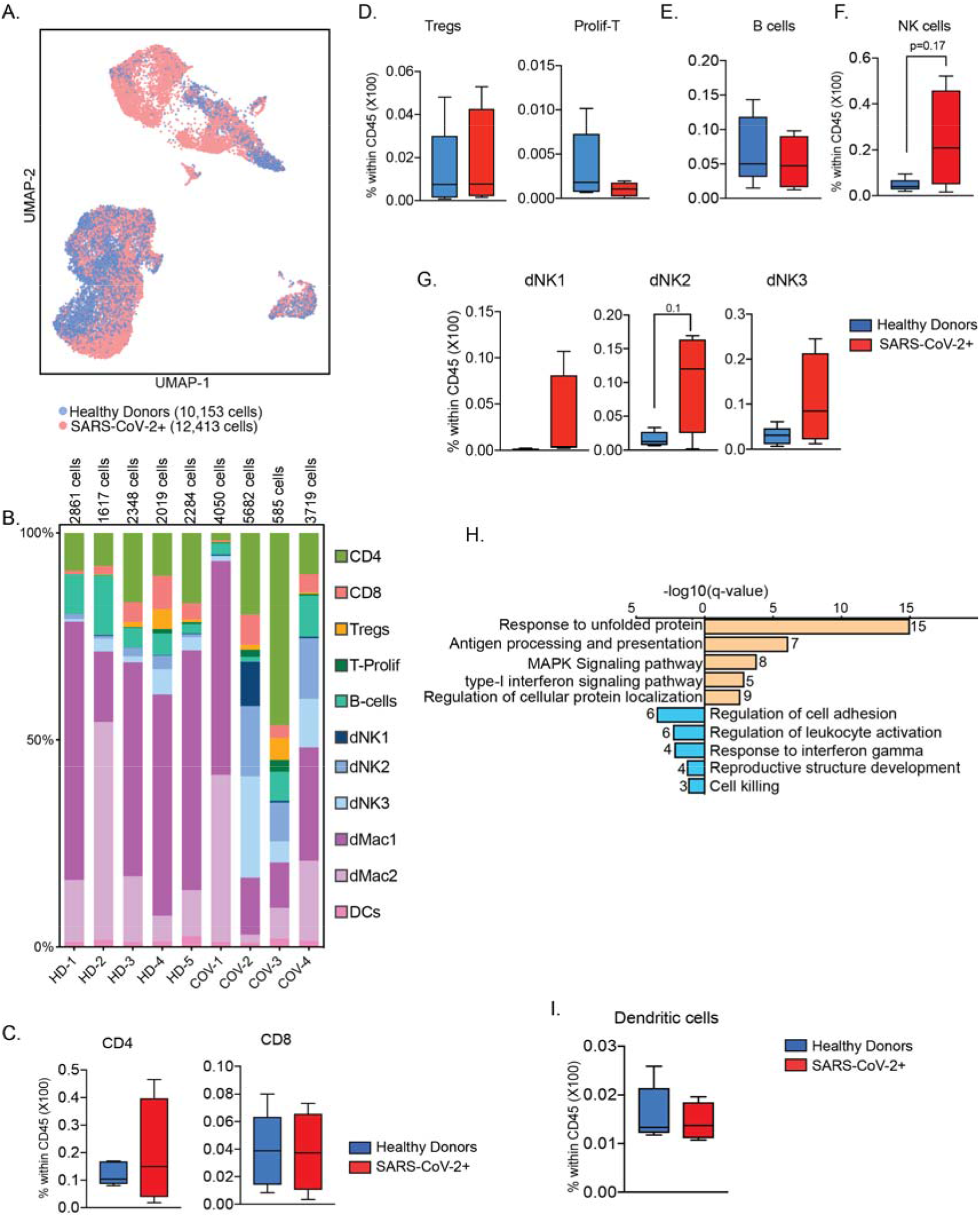
Unsupervised analysis of CD45+ compartment within term maternal decidua and adaptations with asymptomatic SARS-CoV-2 infection. (A) UMAP of term maternal decidual CD45 compartment highlighting cells from healthy donors (blue, 10,153 cells) and mothers with asymptomatic/mild SARS-CoV-2 infection (red, 12,413 cells). (B) Stacked bar graphs comparing per individual immune cell clusters within healthy donors and individuals with infection. (C-G) Box and whiskers comparing frequencies of (C) CD4 and CD8 T cells, (D) regulatory and proliferating T cells, (E) B cells, (F) NK cells, and (G) NK cell subsets within term decidua of healthy donors and mothers with asymptomatic SARS-CoV-2 infection. (H) Functional enrichment of up- (orange) and down-regulated (blue) genes in total decidual NK cells following asymptomatic SARS-CoV-2 infection. Y-axis represents q-values (negative log10 transformed). (I) DC frequencies from single cell clusters

**Supp Figure 4:**
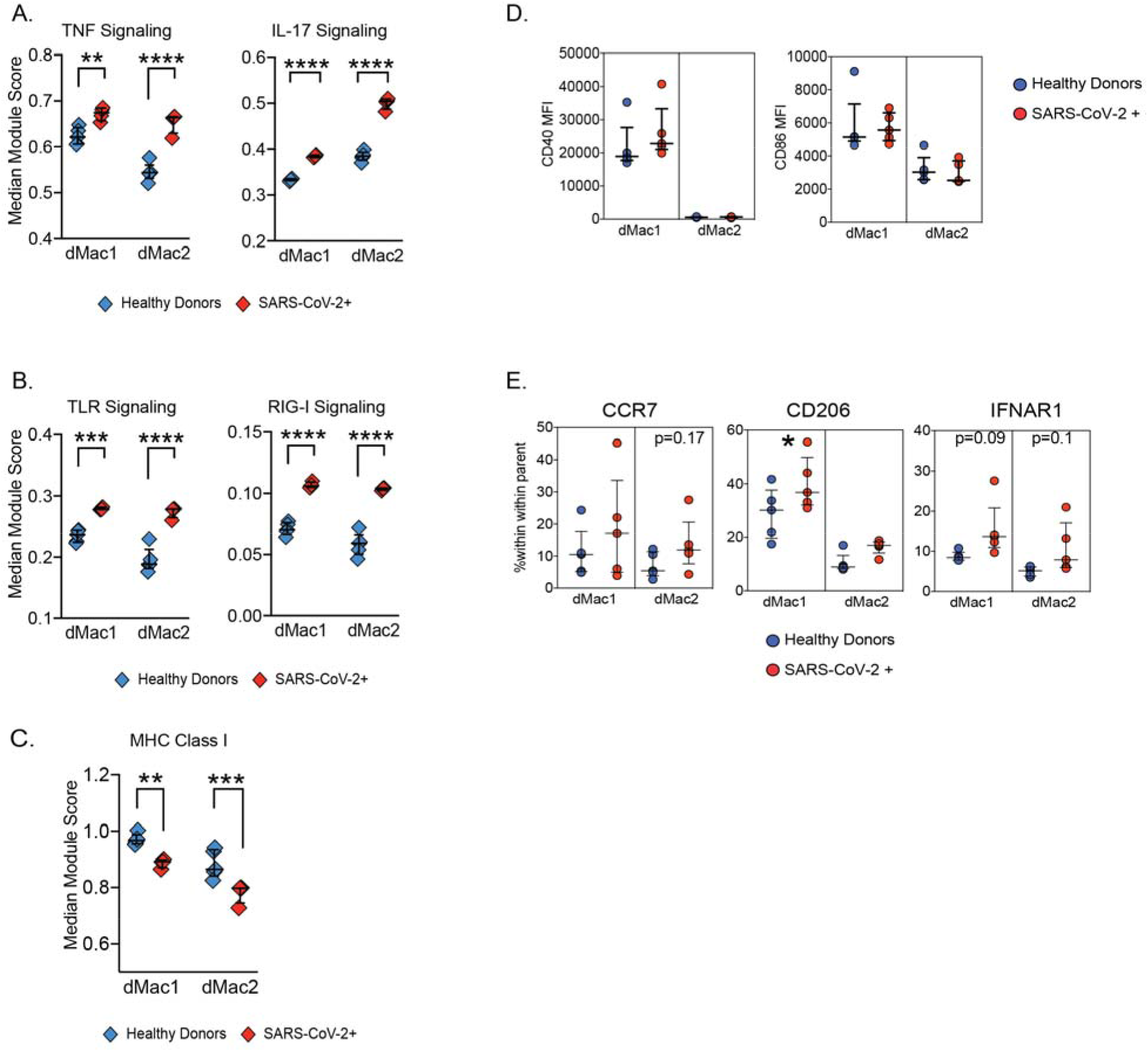
Impact of asymptomatic SARS-CoV-2 infection on term decidual macrophages. (A-C) Dot plots comparing changes in module scores of (A) cytokine signaling pathways (TNF and IL-17 signaling), (B) pathogen sensing pathways (TLR and RIG-I), and (C) MHC Class I pathway in decidual macrophage subsets in healthy donors and mothers with asymptomatic infection. Each dot represents the median module score for the individual. (D) Dot plots comparing median fluorescence of activation markers CD86 and CD40 in decidual macrophage subsets following asymptomatic SARS-CoV-2 infection. (E) Dot plots comparing frequencies of decidual macrophages expressing canonical M1 and M2-like macrophage associated markers (CCR7 and CD206) and interferon receptor (IFNAR1) with asymptomatic SARS-CoV-2 infection. Two group differences were tested using unpaired t-test with Welch’s correction. Error bars represent median values and interquartile ranges. (p-values: ** - p<0.01; *** - p<0.001; **** - p<0.0001)

**Supp Figure 5:**
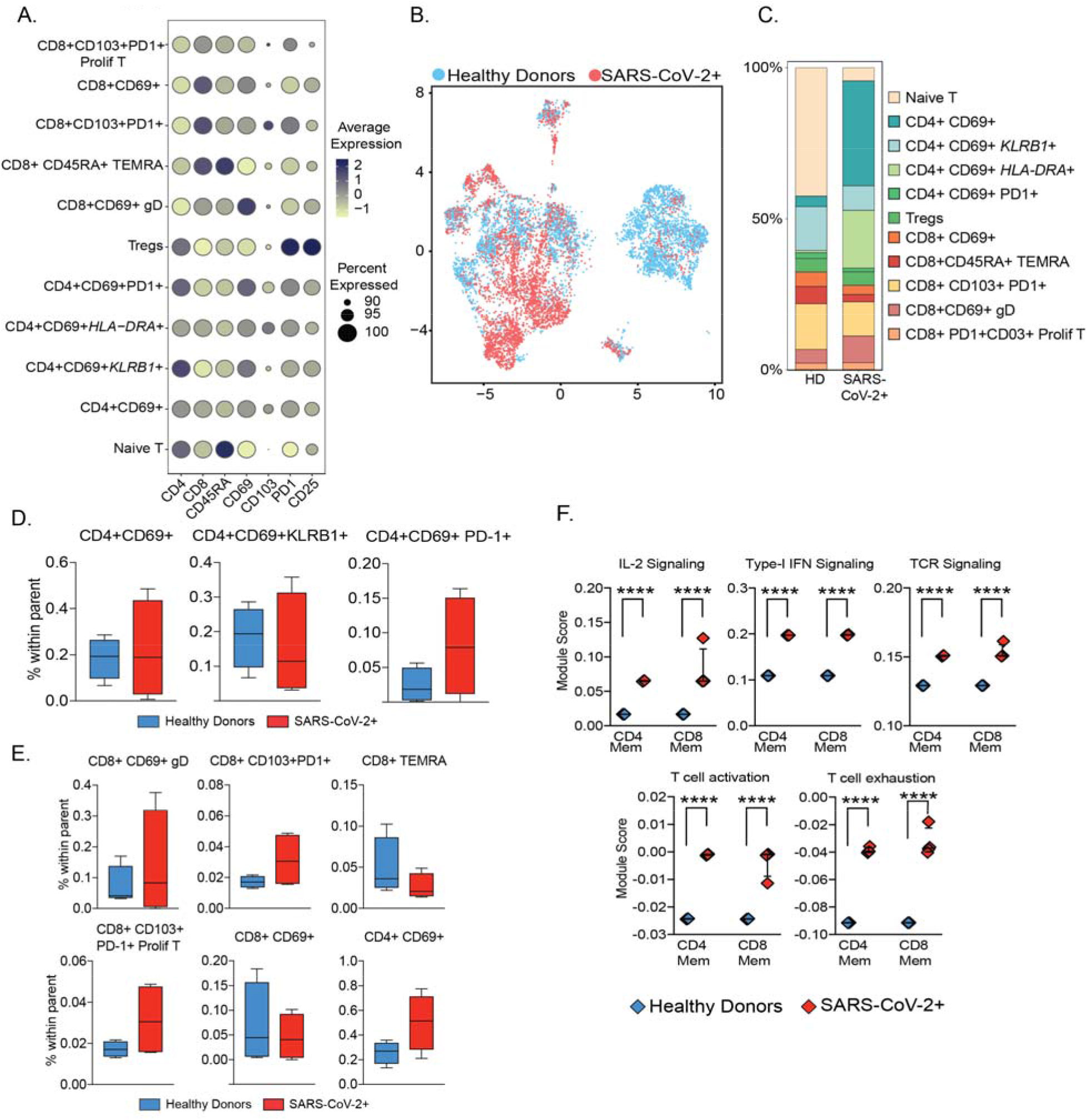
Asymptomatic SARS-CoV-2 infection and T cells at the maternal fetal interface. (A) Bubble plot of cell surface protein expression of markers within each decidua T cell cluster. Size of the bubble represents percentage of cells within each cluster expressing the corresponding marker, whereas color demonstrates magnitude of average surface expression within each cluster. (B) UMAP of term maternal decidual T cell compartment highlighting cells from healthy donors (blue) and mothers with asymptomatic SARS-CoV-2 infection (red). (C) Stacked bar graphs comparing T cell subset proportions between healthy donors and mothers with asymptomatic SARS-CoV-2 infection. (D-E) Box and whiskers comparing (D) memory CD4 and (E) memory CD8 T cell cluster frequencies. (F) Dot plots comparing changes in module scores of T cell signaling pathways in healthy donors and mothers with asymptomatic infection (n=4/group). Each dot represents the median module score for the individual.

**Supp Figure 6:**
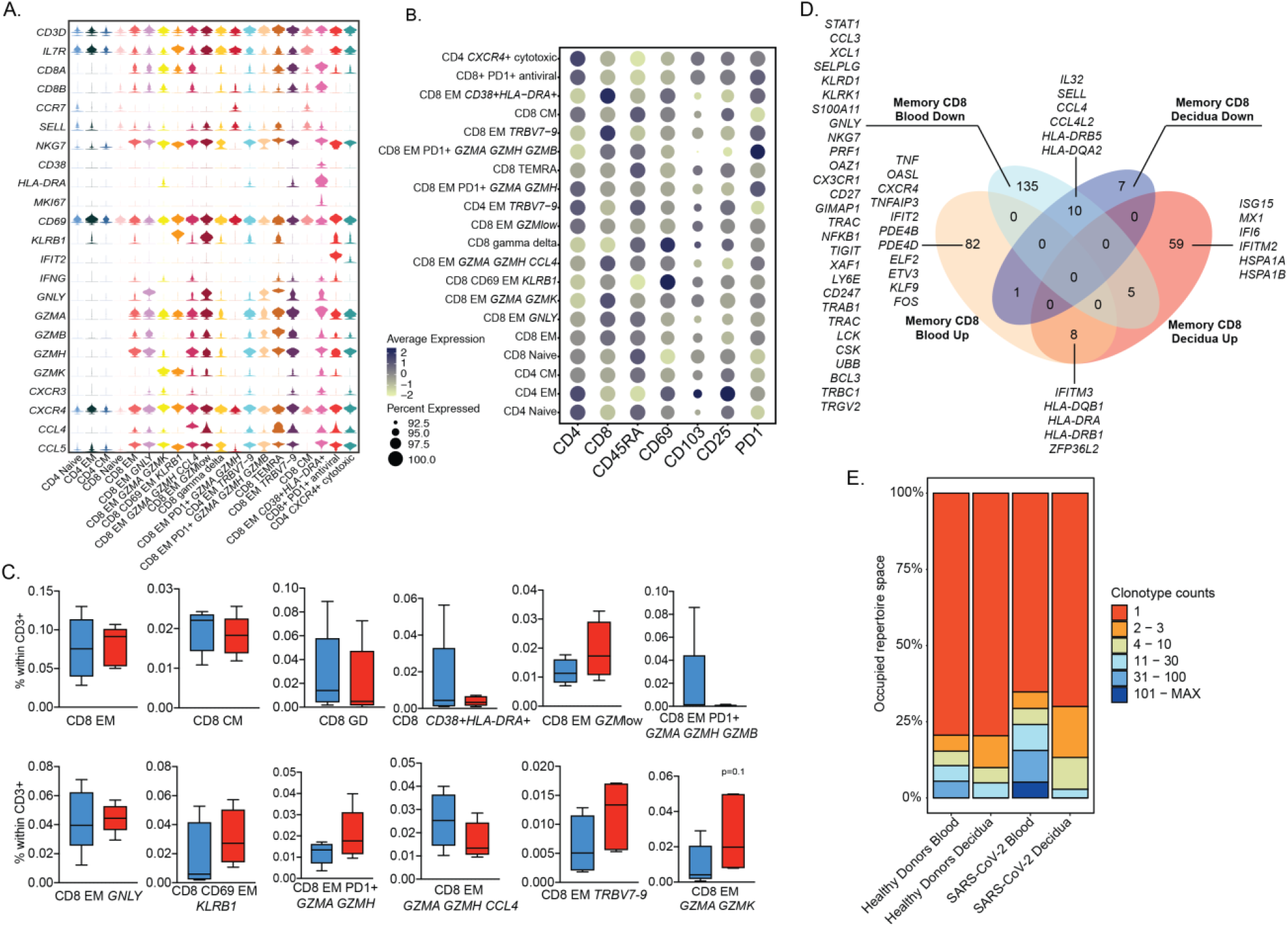
T cell responses to asymptomatic SARS-CoV-2 infection in blood vs. decidua. (A) Stacked violin plots highlighting key gene expression markers of blood T cell subsets within term maternal decidua. Y-axis represents the normalized transcript counts. (B) Bubble plots of cell surface protein expression of markers within each blood T cell cluster. Size of the bubble represents percentage of cells within each cluster expressing the corresponding marker, whereas color demonstrates magnitude of average surface expression within each cluster. (C) Box and whiskers comparing frequencies of CD8 memory subsets in blood with asymptomatic SARS-CoV-2 infection during pregnancy. (D) Venn diagram comparing differentially expressed genes (log2 fold change ≥ 0.4, up or down-regulated, q-value ≤ 0.05) in memory CD8 T cells from blood and decidua. Select genes within this comparison are highlighted. (E) Stacked bar graphs comparing blood T cell clonal sizes with asymptomatic SARS-CoV-2 infection within the occupied repertoire space.

## Acknowledgements

We are grateful to all participants in this study. We thank the MFM Research Unit at OHSU for sample collection. We thank Dr. Jennifer Atwood for assistance with sorting at the flow cytometry core at the Institute for Immunology, UCI. We thank the UCI Genomics and High-Throughput Facility for assistance with 10X library sequencing. Aspects of experimental design figures were generated using graphics from Biorender.com.

## Author Contributions

S.S., N.E.M, and I.M. conceived and designed the experiments. M.R, D.T., and R.E., procured and coordinated specimen collection. S.S., M.Z., B.D., and A.J. performed the experiments. S.S., M.Z., and B.D analyzed the data. S.S., M.Z., and I.M. wrote the paper. All authors have read and approved the final draft of the manuscript.

## Data availability

The datasets supporting the conclusions of this article are available on NCBI’s Sequence Read Archive (SRA# pending).

